# Paired metagenomics and metatranscriptomics reveal metabolic capabilities of uncultivated *Verrucomicrobia* and other bacteria in the Hawaiian bobtail squid reproductive symbiosis

**DOI:** 10.64898/2026.01.08.698421

**Authors:** Andrea M. Suria, Spencer V. Nyholm

## Abstract

Host-associated microbiomes often consist of complex bacterial consortia, many of whose members remain uncultivated and/or have poorly characterized functions. In this study, we used metagenomics and metatranscriptomics to better understand the reproductive defensive symbiosis of the accessory nidamental gland (ANG) of the Hawaiian bobtail squid, *Euprymna scolopes*. We recovered 23 high- and medium-quality metagenome-assembled genomes (MAGs) from the four major ANG symbiont taxa (*Alphaproteobacteria*, *Verrucomicrobia*, *Gammaproteobacteria*, and *Flavobacteriia*) that dominate the *E. scolopes* ANG community. Three *Puniceicoccaceae* MAGs represent the first *Verrucomicrobia* genomes from a cephalopod ANG and are potentially novel species in *Verrucomicrobia* subdivision four. These *Verrucomicrobia* encode the highest diversity of carbohydrate-active enzymes (CAZymes) among the analyzed strains. Metatranscriptomes revealed no differential expression between the ANG and eggs, indicating metabolic stability during symbiont transfer from the ANG tubules to egg jelly coats. Community-wide expression of glycoside hydrolases may enable shared degradation of host O- and N-glycosylated mucins. Genes often important in host-association, such as motility, chemotaxis, and quorum sensing, were broadly expressed amongst the major taxa. Expression of diverse secondary metabolite clusters (e.g., NRPS, PKS, bacteriocins) and competitive mechanisms, like the T6SS, were also expressed and may contribute to mediating competition within the ANG or in egg defense. Overall, this work reveals the genomic and transcriptomic repertoire of the *E. scolopes* ANG microbiome and provides insight on members of the marine *Verrucomicrobia,* a group with growing recognition as important in symbiotic associations.

**Significance:** The *Verrucomicrobia* are a ubiquitous group of growing interest, particularly due to the role of *Akkermansia* in the human gut. However, little is known about aquatic *Verrucomicrobia,* especially those that switch between a free-living and host associated lifestyle. This study furthers our understanding of potential mucin degradation among novel *Puniceicoccaceae* of the *E. scolopes* ANG reproductive symbiosis. Shared expression of carbohydrate utilization pathways among ANG symbionts may reveal networks of competition and cooperation that parallel the complex networks found in the human gut. As well, this study lays the groundwork for further understanding the mechanisms of bacteria-mediated host egg defense, serving as a model for study of other marine defensive symbioses.

## Introduction

In defensive host-microbe symbioses, microorganisms protect their hosts from infection and predation by many diverse mechanisms. While some symbionts produce bioactive compounds that repel harmful organisms, others benefit the host through indirect methods, such as competitive exclusion or bolstering the host’s immune system (reviewed in (1)). For many of these symbioses, the mechanism of defense is often unknown or not well characterized. As well, it is often unclear what benefits the symbionts may gain from the host (2). Uncovering the basis of these interactions can be especially challenging when many symbionts are uncultured and methods to raise gnotobiotic hosts are still in development.

The accessory nidamental gland (ANG) of the Hawaiian bobtail squid, *Euprymna scolopes*, contains one such symbiosis where studying the mechanisms of host defense is complicated by many uncultivated bacteria and host-rearing challenges. The ANG is a female reproductive gland found in many cephalopods that houses diverse bacterial communities (3). In *E. scolopes,* the bacteria are maintained in epithelium-lined tubules **(Figure 1)** and deposited into the jelly coat layer of eggs, where they prevent biofouling and increase juvenile hatch rates (4). The *E. scolopes* ANG community is predominantly composed of *Alphaproteobacteria*, *Verrucomicrobia*, *Gammaproteobacteria,* and *Flavobacteriia* (5, 6). Despite this diversity, the majority of cultured isolates are *Leisingera* and *Ruegeria*, genera from the most dominant taxa in adult squid, the *Alphaproteobacteria* (4, 5, 7, 8). A large diversity of taxa have still evaded culturing to date and little is known about their roles in the community. Notably, there are currently no cultured isolates from the *Verrucomicrobia*, the most abundant community member in immature squid and the second most abundant in adults (9). Manipulating the host for functional studies is complicated further by the fact that the ANG does not develop until a month after hatching, and only if a currently unknown environmental cue is present (10). These factors have necessitated ‘omics and chemical approaches to *in vivo* studies of ANG bacterial metabolism.

**Figure 1.**
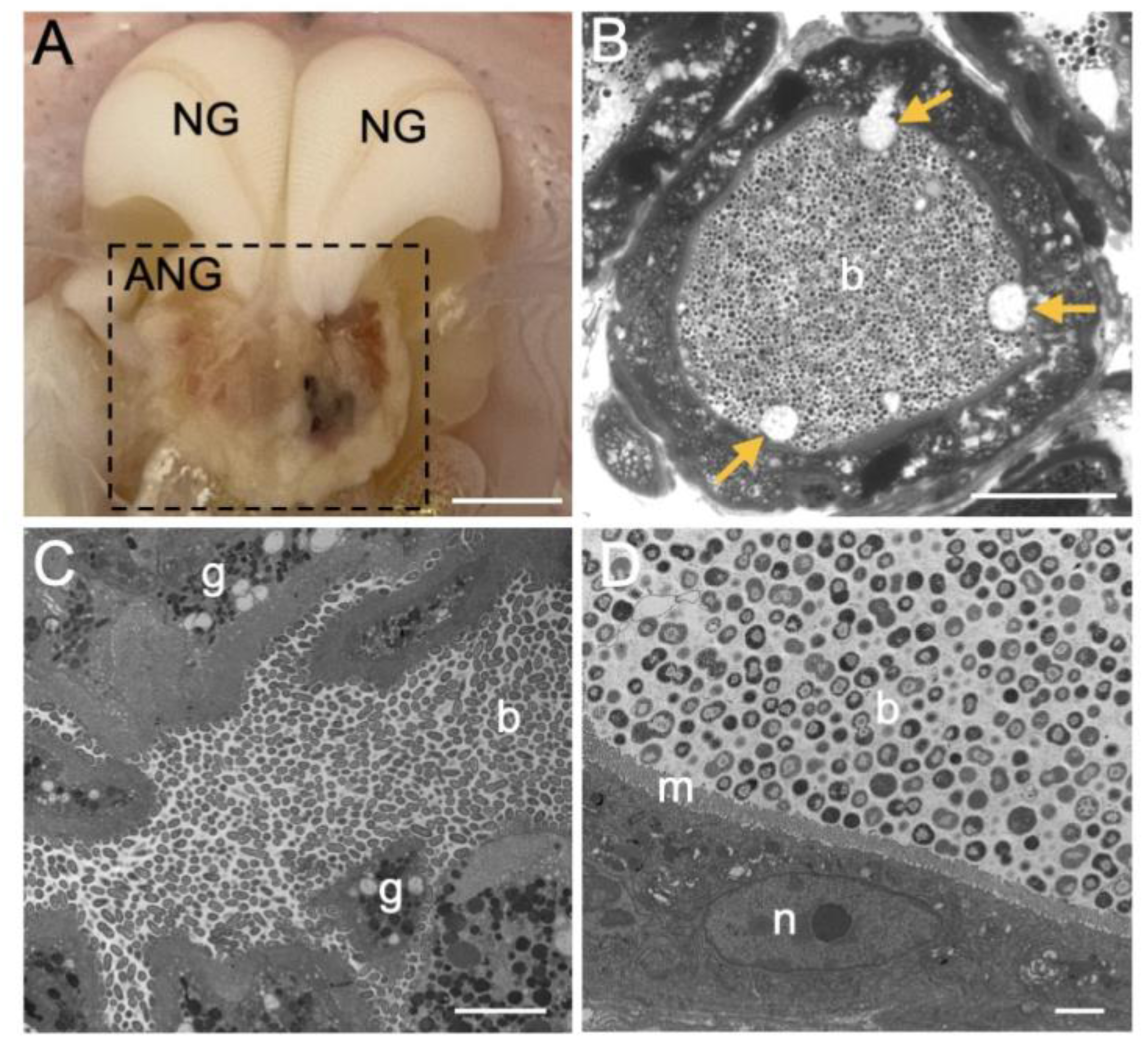
Anatomy of *Euprymna scolopes* accessory nidamental gland (ANG). **(A)** The ANG (boxed) is located beneath the nidamental glands (NG) and is composed of many different colored tubules containing the symbiotic bacteria. Scale: 5 mm. **(B)** Light micrograph cross-section of a tubule shows the bacteria in the lumen and the host epithelium releasing secretory vesicles (yellow arrows) into the lumen. Scale: 40 µm. TEM cross-sections of tubules show bacillus-shaped bacteria **(C)** and coccoid-shaped bacteria **(D)**. Letters: b, bacteria; g, host secretory granules; m, host microvilli; n, host nuclei. Scales: C, 5 µm; D, 2 µm.

While some *in situ* chemical analyses have been performed on *E. scolopes* eggs, little is known about bacterial metabolism in the ANG or eggs. Although the eggs contain a lower bacterial cell density than the ANG, the bacteria do appear to be metabolically active over the course of egg development. Evidence of cell division and the presence of microcolonies have been detected in the eggs (6). A previous LC/MS-MS study of the eggs detected thousands of bacterial metabolites, the majority of which were unclassified (4). To date, there have been no exhaustive *in situ* studies of bacterial metabolites in the ANG, although presence of homoserine lactones has been detected (11). Culture-based *in vitro* assays have identified two antimicrobial metabolites that may play a role in egg defense: indigoidine from *Leisingera* sp. JC1 (7) and bromoalterochromides from *Pseudoalteromonas* sp. JC28 (8). As well, genomic studies have described the primary metabolic potential of many ANG *Alphaproteobacteria* strains, but potential carbon utilization and energy source pathways have not been explored (7, 11).

The pairing of metagenomic and metatranscriptomic sequencing has created a clearer picture of symbiont metabolism for many difficult-to-manipulate symbioses, including the human gut (12), honeybee gut (13), and termite hindgut (14), among others. A paired ‘omics strategy was applied in this study to gain a better understanding of how the *E. scolopes* reproductive symbionts function in both adult hosts and their eggs. Here, we generated metagenome-assembled genomes (MAGs) of many currently uncultured ANG bacteria, including three *Verrucomicrobia* MAGs, and compared the metabolic potential of the four major taxa. Carbon utilization pathways were examined to better understand how the community is maintained and what nutrients the host may provision to its symbionts. A shared mucin degradation pathway was expressed amongst all four taxa, supporting evidence that host mucins may play integral roles in maintaining symbioses. Examining community ANG gene expression also revealed a broader secondary metabolite biosynthetic potential than has been previously observed from isolate genomes. By incorporating these different methods, we present a more comprehensive picture of host-symbiont interactions and lay the groundwork for further studies into the defensive mechanisms of the ANG symbiosis. These findings may guide understanding host-associated metabolism in other complex, environmentally transmitted defensive symbioses.

## Materials and Methods

### Hawaiian bobtail squid sample collection

Hawaiian bobtail squid were collected from Maunalua Bay, Oahu, Hawaii (21°16’51.6“N, 157°43’44.4”W) from 2016 to 2018 **(Table S1)** and shipped to the University of Connecticut. Squid were maintained in individual tanks with artificial seawater for at least one week prior to mating and eggs were collected within 12 h of laying. The jelly coats (JCs) of thirty eggs from the same clutch were pooled as one JC sample per female. Nine sexually mature females were euthanized following protocols approved by the University of Connecticut Institutional Animal Care and Use Committee (A18-029 and A22-004) to sample the ANG. Dissection tools were treated with RNase Away (Ambion) for 1 h prior to use. All samples were surface sterilized with 95% ethanol and washed in filter-sterilized squid Ringer’s solution (FSSR, 530 mM NaCl, 25 mM MgCl_2_, 10 mM CaCl_2_, 20 mM HEPES, pH 7.5) after removal. Samples were either flash-frozen in liquid nitrogen (metagenomics) or placed in Invitrogen RNAlater (metatranscriptomics) and stored at −80°C prior to extractions. TEMs and light micrographs of ANG tissue were prepared as described in (5).

### ANG bacterial DNA extraction and metagenome sequencing

ANG tissue was homogenized in 0.5 ml FSSR with a ground-glass homogenizer and centrifugation was used to separate squid tissue from bacterial cells. DNA was extracted from the bacterial pellet using the MasterPure Complete DNA and RNA Purification Kit (Epicentre) with additional bead-beating. Metagenomic libraries were prepared using the Illumina TruSeq Nano DNA library prep kit and sequenced using the Illumina MiSeq reagent Nano kit v2 (2×250 cycles) for sample PD13 and the Illumina MiSeq reagent kit v2 (2×250 cycles) for all other samples **(Table S2)**. Raw reads were quality filtered, trimmed, assembled, and annotated following methods outlined in **Text S1** and **Figure S1**. Metagenome assembly statistics can be found in **Table S2** and read taxonomy in **Figure S2A**.

### Metagenome Assembled Genome (MAG) binning, phylogeny, and annotation

Individual metagenome assemblies were manually binned into MAGs using Anvi’o v6.2 (15) (see **Text S1**). In accordance with MIMAG recommendations (16) a high quality MAG had 90% genome completion and <10% contamination, while a medium quality bin had 60-89% completion and <20% contamination. Phylogenetic trees were built using GToTree v1.7.06 (17). The genomes of five previous ANG isolates (4) were sequenced using previous methods (8) to include as references. These isolates are *Allomuricauda* sp. ANG21, *Leisingera* sp. JC11, *Roseibium algicola* ANG18, *Ruegeria* sp. ANG10, and *Vibrio* sp. JC34. All MAGs and genomes are uploaded to Genbank under BioProject accession PRJNA940485.

The average nucleotide identity (ANI) of all MAGs was calculated using fastANI (18). An ANI value of >95% was considered to be the same species (19). MAGs were annotated with Prokka v.1.12 (20) using default settings and carbohydrate active enzymes were identified using the dbCAN meta server (21), only retaining hits assigned from two or more tools. BlastKOALA (22) was used to assign KEGG numbers to the predicted proteins. MAGs were screened for secondary metabolite biosynthetic genes using antiSMASH v.3.0 (23).

### ANG and egg jelly coat (JC) RNA extractions

ANGs were homogenized via bead-beating in 1.0 ml of TRIzol Reagent (Invitrogen). Due to difficulty homogenizing egg JCs, frozen samples were first lyophilized using a Labconco FreeZone Plus cascade benchtop freeze dryer for 24 h at −84°C. Lyophilized JCs were then homogenized in 1.0 ml of TRIzol. RNA was extracted following the manufacturer protocol, and treated with TURBO DNase. For ANG samples, eukaryotic host mRNA was selected using the NEBNext Poly(A) mRNA Magnetic Isolation Module (New England BioLabs). Washes from the magnetic beads were pooled and cleaned using the Zymo RNA Clean and Concentrator Kit to collect total bacterial RNA. Bacterial RNA was treated with a 50:50 mixture of Illumina RiboZero bacteria and RiboZero Gold epidemiology rRNA removal probes to remove both host and bacterial rRNA. Since the embryo was removed from all eggs prior to extraction, JC samples were not expected to contain abundant eukaryotic RNA and Poly(A) mRNA removal was not performed. Due to low RNA concentrations, as measured by the Qubit RNA HS assay kit (**Table S3**), JC samples were not treated with rRNA removal probes to avoid sample loss. RNA quality was checked using an Agilent TapeStation 2200.

### ANG and JC Metatranscriptome sequencing and analysis

Metatranscriptomic libraries were prepared using the Illumina TruSeq Stranded mRNA Library Prep Kit. Due to low RNA concentrations, all samples were concentrated to 5 μl using a ThermoSci Savant DNA 120 SpeedVac at 22°C and combined with 13 μl of the “Fragment, Prime, Finish Mix” before proceeding with manufacturer protocol. Libraries were sequenced using the Illumina NextSeq 500 mid-output reagent kit v2 (2×150 cycles) at the Center for Genome Innovation, University of Connecticut.

Complete bioinformatic analyses are detailed in **Text S1** and **Figure S1**. A de novo metatranscriptome was assembled using Trinity v.2.6.6 (24) and assembly statistics can be found in **Table S3**. Eukaryotic mRNA reads were aligned to the assembly using Bowtie2 v.2.3.1 (25). Prokaryotic mRNA reads were either concurrently aligned to reference genomes using BBsplit (26) **(Figure S3, Table S4)** or reads were binned into the four most dominant bacterial taxa, using DIAMOND v.0.9.19 (27) and MEGAN6 (28), and then mapped to the metatranscriptomic assembly **(Figure S2B,C)**. Raw read counts were obtained using featureCounts (29) and TMM normalized in EdgeR v.3.24.3 (30). Differential expression analysis was performed with NOISeq v.2.26.1 (31) with significance cut-offs of p-value **≤** 0.05 and fold change **≥** 2.

## Results and Discussion

Recovery of Metagenome-Assembled Genomes (MAGs) from *E. scolopes* ANG yields uncultured strains and novel *Verrucomicrobia*.

ANG metagenome contigs were manually curated into 94 bins, from which 9 high quality and 14 medium quality MAGs were recovered **(Figure 2, Table S5)**. The remaining 71 bins contained low quality MAGs that were excluded from further analyses. The 23 MAGs kept for analysis all had taxonomic assignments to one of the four major bacterial taxa previously found to make up the core *E. scolopes* ANG community (6): *Alphaproteobacteria* (65% of MAGs), *Flavobacteriia* (17% of MAGs), *Verrucomicrobia* (13% of MAGs), and *Gammaproteobacteria* (4% of MAGs). Of these, two MAG pairs are likely the same species: *Ruegeria* sp. E08Bin6-1/*Ruegeria* sp. E16Bin7-6-1 (97.3% ANI) and *Ruegeria* sp. E08Bin5-7/*Ruegeria* sp. E07Bin8-1 (95.2% ANI) **(Table S6)**. The E08Bin6-1/E16Bin7-6-1 pair were the only MAGs with high similarity (97.3% ANI, **Table S6**) to any existing ANG isolate genome, *Ruegeria* sp. ANG-S4. Similarity of strains from the ANGs of different females supports the conservation of certain species within this symbiosis. As well, ten MAGs represent genera that have not yet been cultured from *E. scolopes*: *Arenibacterium* (E07Bin7-2-1, E16Bin7-2), *Erythrobacter* (E07Bin11-1), *Kordiimonas* (E07Bin3-1), *Paracoccaceae* (E08Bin-5-1, E16Bin-6-3), *Puniceicoccaceae* (E07Bin2-1, PD13Bin3-1, PD13Bin3-4), and *Rhizobiaceae* (E07Bin6-1).

**Figure 2.**
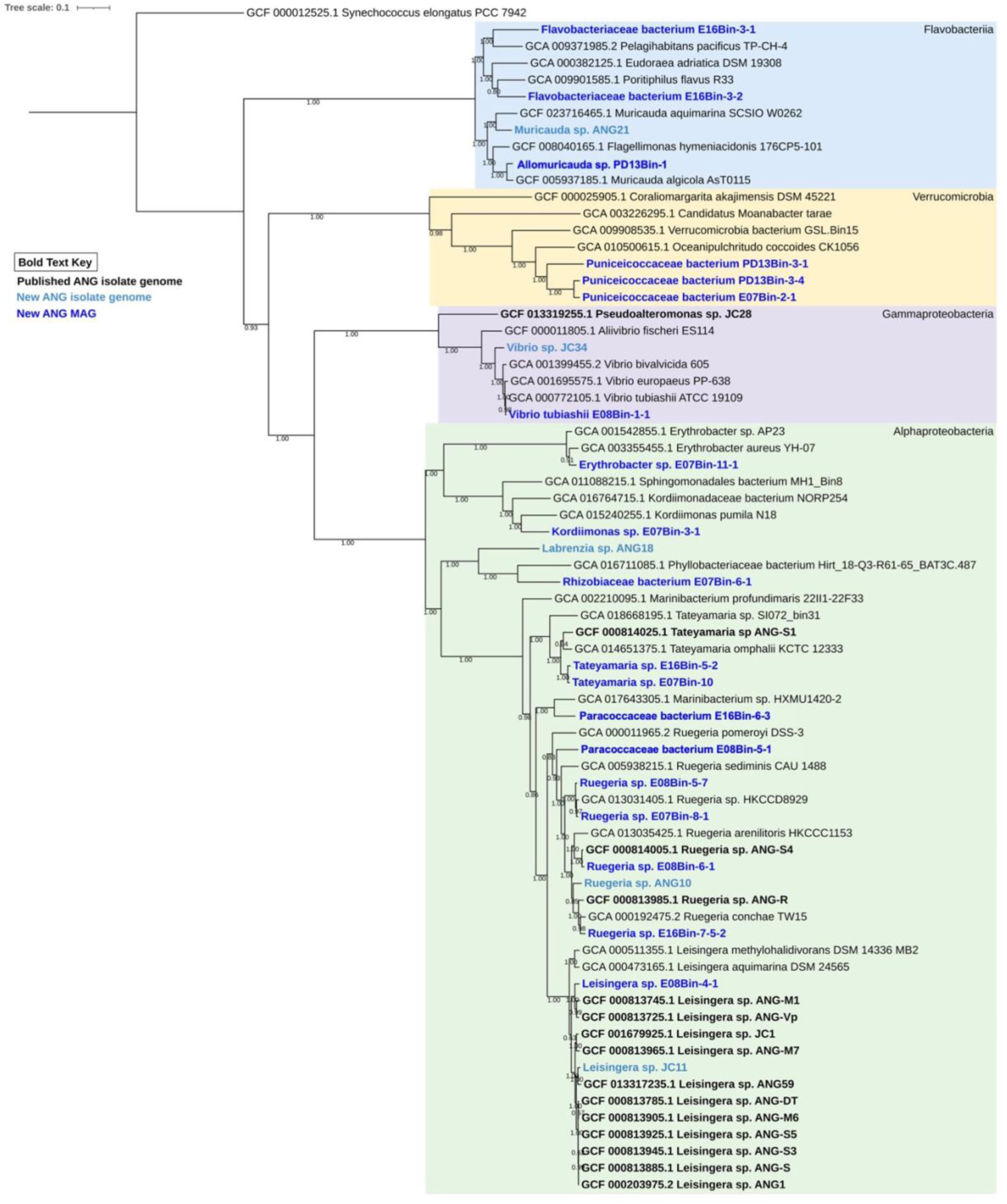
Phylogenetic tree of MAGs and ANG isolate genomes. Metagenome-assembled genomes (MAGs) are in bold dark blue font, newly sequenced ANG isolate genomes are in bold light blue font, and previously published ANG isolate genomes are in bold black font. Colored ranges highlight the four major taxa of the ANG community: *Alphaproteobacteria* (green), *Verrucomicrobia* (yellow), *Gammaproteobacteria* (purple), and *Flavobacteriia* (blue). Bootstrap values are displayed at each node. Tree was constructed using 74 single copy genes (SCG).

The three *Puniceicoccaceae* MAGs are of particular interest since they represent the first *Verrucomicrobia* genomes from an ANG symbiosis. *Puniceicoccaceae* bacteria E07Bin2-1, PD13Bin3-1, and PD13Bin3-4 are all high-quality assemblies belonging to *Verrucomicrobia* subdivision 4 **(Figure S4)**. The closest reference genome, *Oceanipulchritudo coccoides* CK1056, is an environmental isolate from marine sediment (32). These MAGs did not have an ANI value above 70% with any *Verrucomicrobia* reference genomes, and had ANI values <70-86% compared to each other **(Table S6)**, indicating they are novel taxa in this phylum.

### Carbohydrate active genes are predicted in all genomes and widely expressed in *Puniceicoccaceae* MAGs

The MAGs and isolate genomes were screened against the Carbohydrate-Active enZYmes (CAZy) database to determine the potential carbon sources the symbiotic community can utilize. A total of 235 CAZy families were predicted across the 43 genomes analyzed **(Table S7)**. CAZy predictions per genome ranged widely from 0.63% up to 5.37% of the total coding sequences **(Figure 3A)**. GT families were the most broadly distributed with an average of 31 predictions per genome, GT4 being the most abundantly predicted. Family GT4 typically encodes enzymes that utilize GDP-mannose or UDP-GlcNAc as nucleotide-sugar donors and can play roles in lipopolysaccharide, exopolysaccharide, and secondary metabolite synthesis (33).

**Figure 3.**
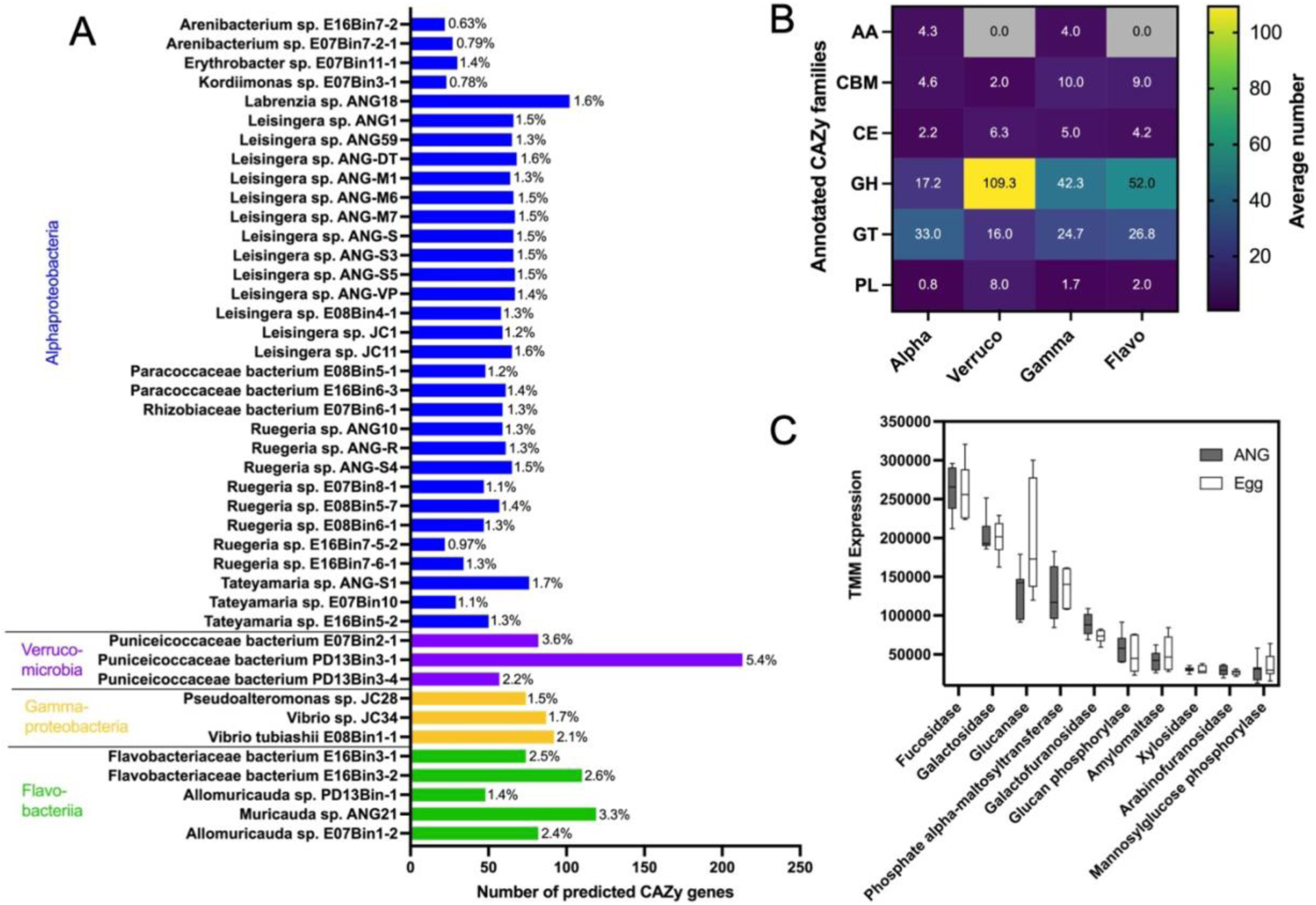
Diversity of carbohydrate active enzymes predicted in all symbiont genomes and expressed in *Puniceicoccaceae* bacterium PD13Bin3-1. **(A)** Number of predicted carbohydrate active enzyme (CAZy) genes in each symbiont MAG or isolate genome. Percentage beside bar indicates the percent of predicted coding sequences that are CAZy predictions. **(B)** Heatmap of the average number of CAZy families predicted in each genome from each of the top four taxa. Families: AA, Auxiliary Activities; CBM, Carbohydrate Binding Module; CE, Carbohydrate Esterases; GH, Glycoside Hydrolases; GT, Glycosyl-Transferases; PL, Polysaccharide Lyases. **(C)** Average TMM normalized expression of the top 10 most highly expressed glycoside hydrolase families in *Puniceicoccaceae* bacterium PD13Bin3-1 in both the ANG and eggs.

The *Verrucomicrobia* MAGs possessed the greatest average number of glycoside hydrolases of all major taxa **(Figure 3B)**. Similar to other marine *Verrucomicrobia* (34), *Puniceicoccaceae* bacterium PD13Bin3-1 had a high number of predicted CAZy genes (212 genes) and the greatest diversity of glycoside hydrolase families (64 families) of all genomes analyzed. Of these, fucosidases, galactosidases, and glucanases were the most highly expressed in both the ANG and eggs **(Figure 3C)**. Expression levels varied amongst the three *Verrucomicrobia* MAGs, with phosphate alpha-maltosyltransferase expression highest in PD13Bin3-4 and N-acetylglucosaminidase expression highest in E07Bin2-1 **(Figure S5, Table S8)**. As discussed further below, these enzymes could play roles in host-mucin degradation.

### Symbionts express similar host-associated gene profiles in both the ANG and eggs

Metatranscriptomes were sequenced from paired ANG and egg samples to determine the symbiont metabolic profiles as they move through these host-associated environments. Although we hypothesized the transition from the mother’s ANG to externally laid eggs might induce a transcriptional response in the bacteria, there were no significantly differentially expressed genes (**Figure S6, Table S9)**. However, distinct metabolic profiles were detected between the four major bacterial taxa, suggesting specialized roles within the community **(Figure 4, Table S10)**. Carbohydrate, amino acid, and nucleotide metabolisms were the highest expressed pathways for all taxa. Highest TMM normalized expression was seen in carbohydrate metabolism of egg *Gammaproteobacteria* and ANG *Verrucomicrobia*. While *Flavobacteriia* had the lowest carbohydrate metabolism expression, they had the highest expression of purine, sphingolipid, and glycan degradation metabolism genes.

**Figure 4.**
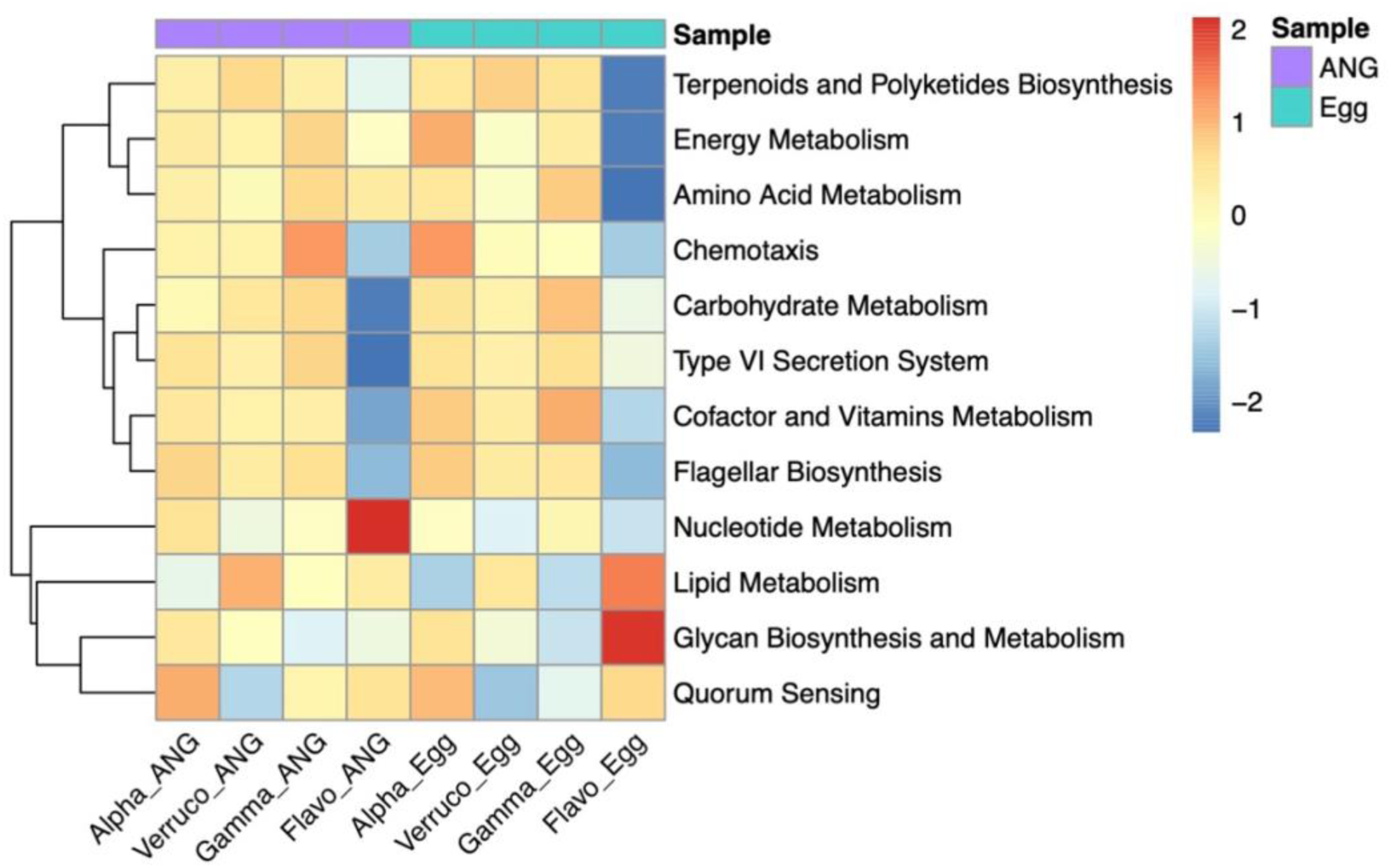
Expression of KEGG metabolic pathways by bacterial symbionts in the ANG and eggs. Metatranscriptome reads sequenced from the ANG (purple samples) and freshly laid eggs (teal samples) were analyzed with the KEGG pathway database. TMM normalized values were z-scaled to compare expression between the four major bacterial taxa: *Alphaproteobacteria* (Alpha), *Verrucomicrobia* (Verruco), *Gammaproteobacteria* (Gamma), and *Flavobacteriia* (Flavo). Hierarchical clustering shows patterns of higher (red) or lower (blue) pathway expression that differed between the taxa.

Lipid biosynthesis is crucial for membrane development, but it can also be critical for host-microbe signaling (35). Glycerophospholipid metabolism was the most highly expressed pathway, with ANG *Flavobacteriia* having higher expression than all other groups, followed by ANG *Verrucomicrobia* **(Figure 4, Table S10)**. The second most highly expressed pathway was sphingolipid metabolism by egg *Flavobacteriia*. Symbiotic sphingolipid-producing bacteria have been shown to maintain host-mucosal homeostasis (36, 37).

Expression of different energy metabolism pathways suggests a mixed use of aerobic and anaerobic respiration in the ANG and eggs. Oxidative phosphorylation was the highest energy metabolism pathway expressed for all groups. Cytochrome c oxidase subunits, *coxABC*, used during aerobic respiration, had higher expression in *Flavobacteriia* and *Gammaproteobacteria*. The *Alphaproteobacteria* expressed the cbb3-type cytochrome c oxidase in both the ANG and eggs, which is usually expressed in low oxygen environments. Both the *Alphaproteobacteria* and *Verrucomicrobia* expressed genes for extracellular nitrate import and nitrate/nitrite reduction to either ammonia or nitric oxide. Thus, the *Alphaproteobacteria* and *Verrucomicrobia* may be adapted to lower oxygen microenvironments, which may occur within ANG tubules and eggs at the center of a clutch.

Exopolysaccharide (EPS) production has been associated with establishing symbioses in squid and legumes (38, 39). All strains expressed some genes required for colanic acid and succinoglycan EPS production, with highest expression in the ANG *Flavobacteriia*. Complete pathway expression was seen for alginate and poly-N-acetyl-glucosamine biofilm formation. Peptidoglycan biosynthesis genes were expressed by all groups, but was highest in both the ANG and egg *Alphaproteobacteria*. This corroborates previous microscopy data to show strains are actively dividing in both of these environments (5, 6).

Motility and chemotaxis can play key roles during host colonization. Both flagellar biosynthesis and chemotaxis genes were expressed in the ANG and eggs by all taxa except the *Flavobacteriia* **(Figure 4)**. Expression of chemoreceptors varied by taxa. The aerotaxis receptor, *aer*, was expressed in *Alphaproteobacteria* and *Gammaproteobacteria*, but not in *Verrucomicrobia* nor *Flavobacteriia*. The *Verrucomicrobia* expressed *tap*, a peptide receptor in both the ANG and eggs. The aspartate and maltose receptor, *tar*, was expressed in all groups except the ANG *Alphaproteobacteria*. The ANG *Gammaproteobacteria* also expressed the ribose and galactose receptor, *trg*. Further research will be needed to determine specific chemoattractants in these environments.

Quorum sensing plays crucial roles in regulating many symbiosis-related genes. Previous studies have shown that several *E. scolopes* ANG isolates possess quorum sensing genes and can produce homoserine lactones in culture (7, 11). Here we observed that the *Alphaproteobacteria* had the highest expression of the *luxIR* homologs, *rhiIR*. Low expression of the acyltransferase, *lsrF*, the repressor, *lsrR*, and the AI-2 synthase, *luxS* was observed in ANG *Gammaproteobacteria*. It remains to be determined which functions are regulated by quorum sensing in these taxa.

The type VI secretion system (T6SS) is a molecular syringe used for toxin delivery during microbial competition and host interactions. T6SS genes were most highly expressed by the *Gammaproteobacteria* and *Alphaproteobacteria,* with low to no expression by the *Flavobacteriia* and *Verrucomicrobia*. In the *E. scolopes* light organ, some *V. fischeri* strains use the T6SS to compete for colonization sites (40). Since the ANG is composed of many different tubules, each dominated by different taxa (5, 41), and many ANG isolates possess T6SS genes (42), we hypothesize that the T6SS may also facilitate competition between strains in the ANG.

### ANG and egg bacteria collectively express all genes necessary to cleave O- and N-glycosylated host mucins

Host mucins play a key role in symbiosis, acting as both a physical barrier for epithelial cells and a dynamic environment for bacterial interactions and metabolism (43–45). A number of mucin biosynthesis genes with amino acid similarity (22-53%) to vertebrate mucins were detected in the *E. scolopes* host ANG transcripts: three MUC3A isoforms, one MUC22 isoform, three MUC24 isoforms, and 13 mucin-like protein isoforms **(Figure S7, Table S11)**. Expression of various transferase enzymes suggest these mucins may be either O- or N-glycosylated, and decorated with either fucose or xylose. Genes for synthesizing core 1, 2, 3, and 4 O-glycosylated mucins were expressed, as well as N-glycan precursor and paucimannose-type N-glycosylated mucin **(Figure S7)**. Though expression levels were relatively low, it should be noted that little is known about mucin biosynthesis genes in cephalopods, and expression may be higher in as yet uncharacterized genes.

The host mucin biosynthesis expression was compared to expression of bacterial mucin degradation genes to determine if the symbiotic community can utilize putative host glycans as an energy source. Based on the combination of host/symbiont expression and what is known about mucin glycosylation in other mollusks (46), hypothetical models of N- and O-glycosylated mucins were created **(Figure 5B)**. The model N-glycan is based on hemocyanin of the snail, *Helix pomatia* (46). Hemocyanin also serves as the oxygen transporter of *E. scolopes* hemolymph. The model O-glycan is based on the hypothetical model presented by Tailford *et al.* (44).

**Figure 5.**
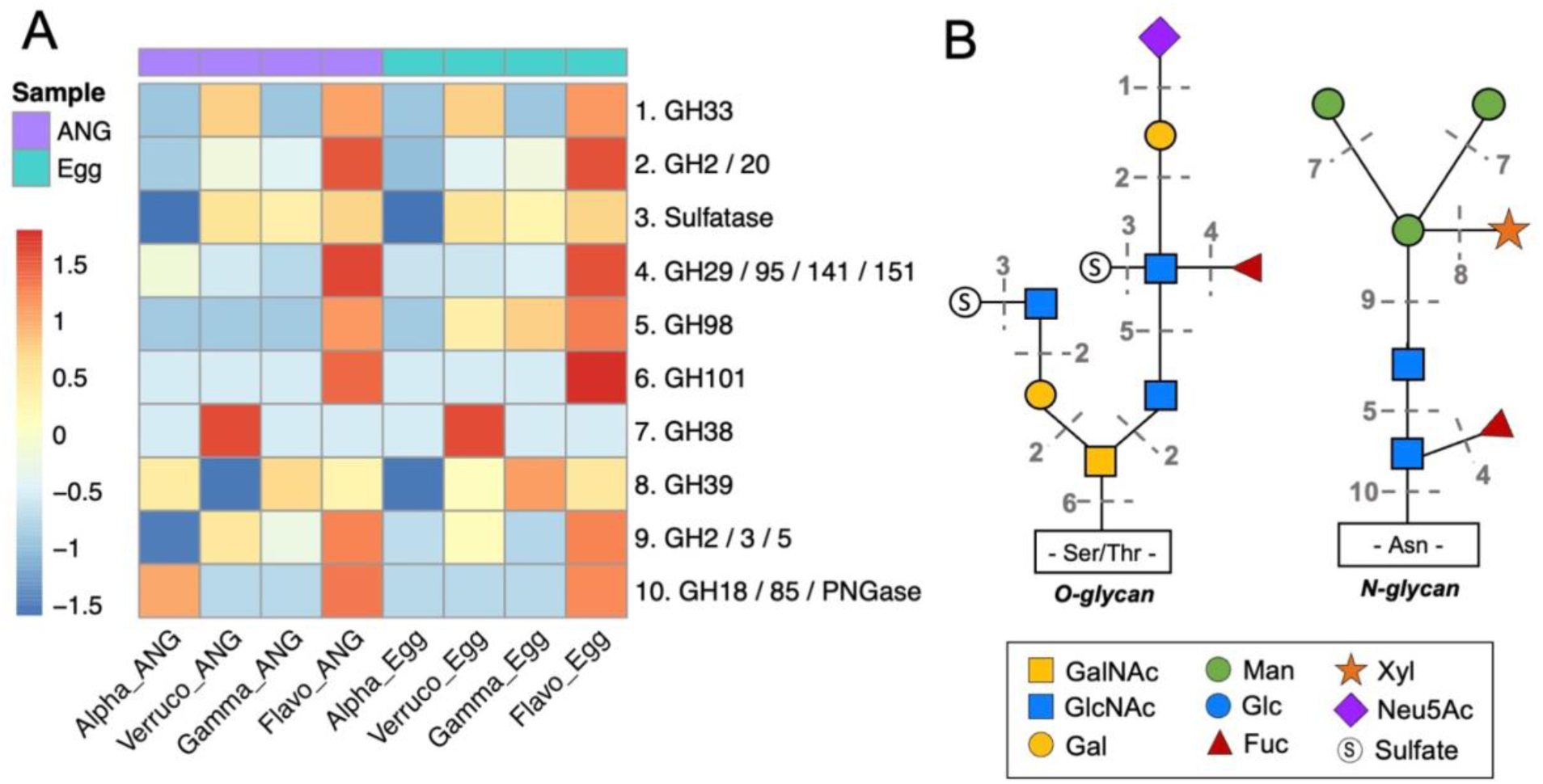
Expression of predicted glycan degradation genes by the ANG and egg bacterial communities. **(A)** Expression of predicted glycan degradation genes in the ANG (purple samples) and eggs (teal samples) was TMM normalized and z-scaled for comparison among the four major taxa: *Alphaproteobacteria* (Alpha), *Verrucomicrobia* (Verruco), *Gammaproteobacteria* (Gamma), and *Flavobacteriia* (Flavo). Glycoside hydrolase (GH) families shown: GH33, sialidases; GH2/GH20, beta-galactosidases; GH29/GH95/GH141/GH151, fucosidases; GH98, galactosidases; GH101, N-acetylgalactosaminidases; GH38, alpha-mannosidases; GH39, xylosidases; GH2/GH3/GH5, beta-mannosidases; GH18 / GH85, peptide:N-glycosidase. **(B)** Hypothetical O-glycosylated and N-glycosylated mucin models. Numbered dotted lines correspond to the enzymes in the heatmap, indicating where each enzyme may cleave off a mucin residue. Legend abbreviations: GalNAc, N-acetylgalactosamine; GlcNAc, N-acetylglucosamine; Gal, galactose; Man, manose; Glc, glucose; Fuc, fucose; Xyl, xylose; Neu5Ac, N-acetylneuraminic acid.

Bacterial expression of ten glycoside hydrolase families and sulfatases were detected **(Figure 5A)**, which may be involved in cleaving residues from these mucins. While no singular bacterial taxa expressed all genes to cleave all residues from either hypothetical glycan, the community as a whole expressed all necessary degradation genes **(Figure 5A, Table S12)**. The *Flavobacteriia* had the most complete expression of mucin degradation genes, only missing GH38 enzymes. These enzymes are alpha-mannosidases that can cleave the terminal mannose on the hypothetical N-glycan. The *Verrucomicobia*, however, did have the highest expression of GH38 enzymes. The *Gammaproteobacteria* had the highest expression of GH39 enzymes in both the ANG and the eggs, potentially filling this pathway gap for all other taxa. The most highly expressed enzymes for all taxa were the fucosidases (GH29, GH95, GH141, GH151), which can cleave fucose residues, and the beta-galactosidases (GH2 / GH20), which can cleave GalNAc and GlcNAc from the first GalNAc of O-glycans. Summing the TMM values of enzymes required to degrade either O-glycans or N-glycans for all bacteria, there was slightly higher O-glycan degradation genes expressed in both the ANG and eggs (ANG: 62,333 TMM, egg: 58,867 TMM) than N-glycan degradation genes (ANG: 46,656 TMM, egg: 42,412 TMM). Further research is necessary to determine if shared use of these enzymes is sufficient to fully degrade mucins in the ANG and eggs, or if certain strains are specialized to certain degradation steps.

### Diverse secondary metabolite biosynthesis genes are expressed in the ANG and eggs

The ability of *E. scolopes* egg symbionts to inhibit fungal and bacterial infections is hypothesized to be due to secretion of antimicrobial compounds (4). To examine this potential at the community level, antiSMASH was used to predict secondary metabolite biosynthesis gene clusters in the ANG metagenome coassembly. A total of 255 secondary metabolite biosynthetic gene clusters from 16 secondary metabolite classes were predicted **(Figure 6A)**. The largest proportions of predicted clusters were homoserine lactones (25.4%), bacteriocins (17.6%), and terpenes (15.6%). These cluster classes have been previously predicted in ANG isolate genomes (7, 11), as well as in the newly assembled MAGs **(Table S13)**, however several new secondary metabolite classes were predicted from the community using this method. New predictions included acyl amino acid, arylpolyene, cyanobactin, nucleoside, T1-/T3-PKS, and thiopeptide biosynthetic gene clusters.

**Figure 6.**
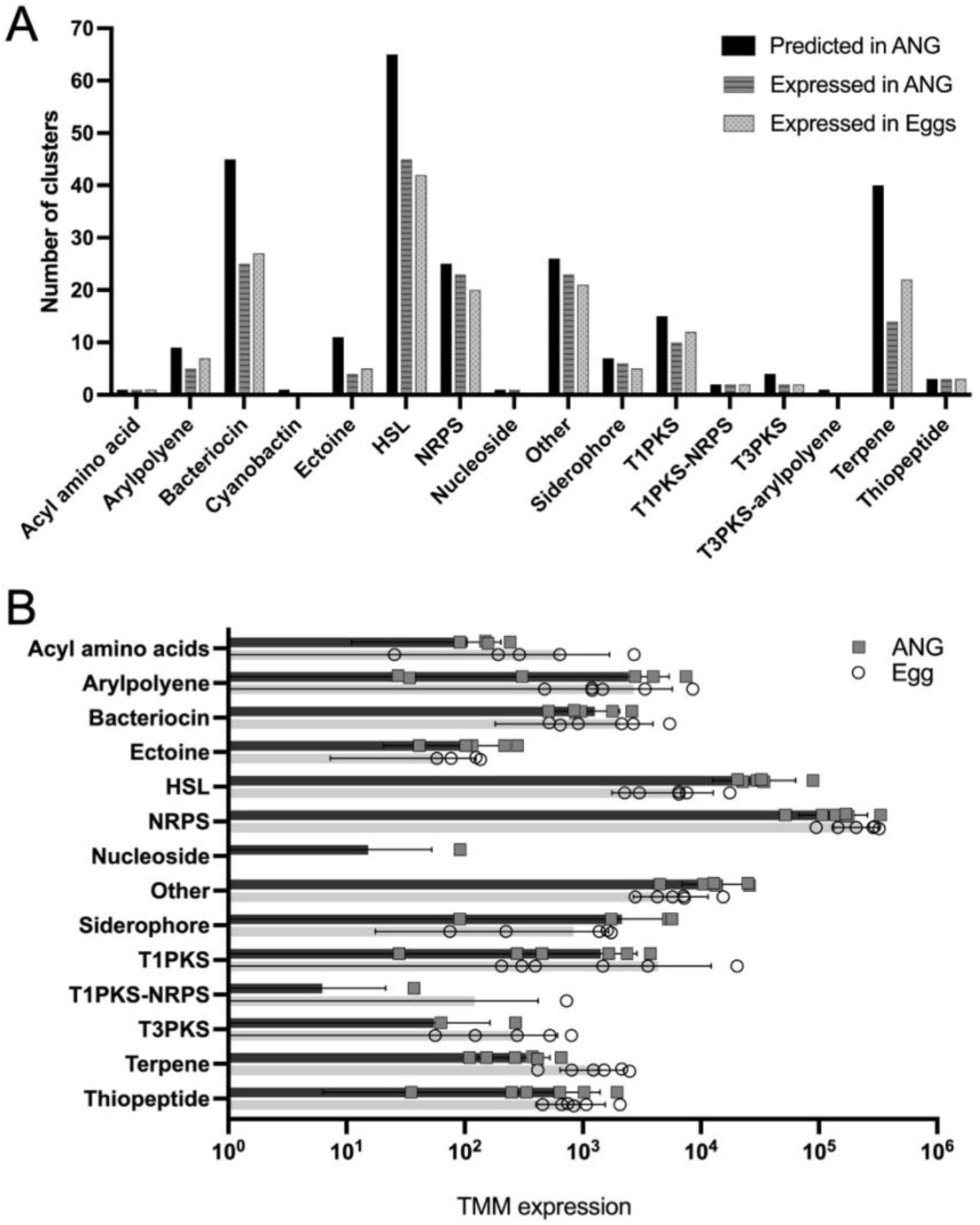
Expression of secondary metabolite gene clusters in the ANG and eggs. **(A)** Number of secondary metabolite gene clusters predicted by antiSMASH in the ANG metagenome (black bars) and clusters expressed in the ANG (dark gray, bars) and eggs (light gray, dots). **(B)** TMM normalized expression levels of the core biosynthetic genes of each predicted cluster in the ANG (black bars, squares) and eggs (gray bars, circles). Symbols represent the expression of biological replicates (n=6). Abbreviations: HSL, homoserine lactone; NRPS, nonribosomal peptide synthetase; T1PKS, type 1 polyketide synthase; T3PKS, type 3 polyketide synthase.

AntiSMASH defines a cluster as genes that match the profile HMMs within <10kb mutual distance but also includes flanking genes up to 20 kb on either side of the last gene hit, which may include accessory genes required for biosynthesis (47). For this reason, not every gene in a predicted gene cluster may be involved in synthesizing the predicted secondary metabolite. Due to this uncertainty, we counted a cluster as expressed if all core biosynthetic genes were expressed, although we acknowledge that expression of accessory genes may be required for complete biosynthesis. Following this rule, 65% (n=163) of the predicted clusters were expressed in the ANG and 66% (n=168) in the eggs. Almost all categories of predicted secondary metabolites were expressed except the cyanobactin and type 3 polyketide-arylpolyene **(Figure 6B)**. For the majority of classes, the number of unique clusters expressed in the ANG and eggs was similar, except in the terpenes, where eight clusters were expressed in the eggs that were not expressed in the ANG **(Table S14)**.

The most highly expressed secondary metabolite class in both the ANG and eggs was the nonribosomal peptide synthetases (NRPS) **(Figure 6B, Table S14)**. NRPS clusters synthesize peptides independent of mRNA and ribosomes, creating a wide diversity of natural products. These compounds are often bioactive and can include antibiotics and siderophores (48), which may play roles in *E. scolopes* egg defense. Another secondary metabolite class that can produce bioactive compounds is the polyketide synthases (PKS). Three PKS groups had higher expression in the eggs: T1PKS (207.4% higher), T3PKS (437.6% higher), and hybrid T1PKS-NRPS (1,851.6% higher). While we could not predict the products of these NRPS or PKS clusters from the metagenomic data alone, this is an area of future research.

Some categories of secondary metabolites may play indirect roles in potential egg defense. Homoserine lactones (HSLs) regulate gene expression via quorum sensing, and may control expression of antimicrobials. The HSLs were the second most highly expressed category after NRPSs. These clusters were expressed 426.5% higher in the ANG than the eggs. This can be expected since bacteria are at a higher cell density in the ANG, suggesting quorum sensing may be more active in the ANG than the eggs. Siderophores were also expressed 155.6% higher in the ANG than in the eggs. By increasing the ability to sequester iron, siderophores may facilitate competitive exclusion to provide host defense in the eggs.

## Conclusions

The paired metagenomic and metatranscriptomic approach used here has provided a more comprehensive view of the bacterial community in the *E. scolopes* ANG symbiosis. Recovery of 23 high and medium quality MAGs has expanded the phylogenetic diversity of available ANG genomes, including novel lineages in the *Verrucomicobia*. The *Verrucomicrobia* remain difficult to culture from most environments due to slow growth, microcolony size, specialized nutrient requirements, and an inability to grow on typical agar-based media (49, 50). Only a few animal-associated *Verrucomicrobia* strains have been cultured to date, from such hosts as ants (51), termites (50), sponges (52), corals (53), pythons (54), and the human gut (55). Similar to these strains, and others associated with brown algae (56), the ANG *Verrucomicrobia* MAGs possess a diverse repertoire of genes that have been known to degrade complex polysaccharides. As CAZy families encompass a broad range of enzymatic activities, confirmation of the exact carbohydrate substrates will depend on future isolate testing coupled with *in vivo* glycomics. Future work will also utilize the CAZy expression profiles generated here to guide *Verrucomicrobia* culturing, as has been successful in culturing the *Rikenella*-like isolate from the leech gut (57).

There remains a strong desire to culture ANG *Verrucomicrobia* isolates in particular to further understanding of ANG development in *E. scolopes*. Previous 16S rRNA gene sequencing of wild-caught bobtail squid ANGs revealed that the *Opitutae* (*Verrucomicrobia* class) were dominant in small females and gradually became succeeded by *Alphaproteobacteria* in larger, mature females (9). As well, development of the ANG in lab-raised squid requires an unknown environmental trigger, of which the *Verrucomicrobia* are hypothesized to play a role (10). Availability of cultured ANG *Verrucomicrobia* isolates may further development of a gnotobiotic ANG symbiosis model.

Glycan degradation genes were among the most highly expressed carbohydrate pathways in ANG *Verrucomicrobia*, which could play a role in tubule colonization patterns. For strain PD13Bin3-1, expression was dominated by fucosidases, which can cleave fucose from complex carbohydrates. This is similar to the human gut-associated *Verrucomicrobia*, *Akkermansia muciniphila*, which can utilize different fucosidases to remove fucose from O-glycosylated mucins (58). The varied expression of glycoside hydrolases among the different major taxa could support the hypothesis that host-provided nutrients influence tubule colonization. Different *E. scolopes* ANG tubules contain either secretory or non-secretory epithelial cells **(Figure 1)** (5), which could deliver different nutrients or possess different types of glycosylated mucins. Fluorescence *in situ* hybridization has shown that different symbiotic taxa are often found in different tubules (5, 41). Provisioning of different nutrients could lead to priority effects during tubule development, as has been seen in the infant gut and plant phyllospheres (59).

Although symbiont partitioning occurs in the ANG, the extant of symbiont interactions within the ANG and eggs is not yet understood. Expression of potential competitive mechanisms in the ANG, such as bacteriocins and T6SSs, may maintain taxa dominance in different tubules. This may be particularly important during egg-laying, when there is potential for symbiont mixing and population depletion as tubule contents are deposited into the egg jelly. The eggs appear to lack strict symbiont partitioning and only possess a membrane-like structure that separates the jelly layers (4, 5). This could facilitate further interactions between symbionts. For example, in the case of glycan degradation, symbionts may benefit from shared resources like secreted glycoside hydrolases to fully utilize the limited mucins available in the egg and potentially allow for cross-feeding, as seen by human gut isolates (60). Mucin degradation has been shown to depend on ecological networks in human gut-associated symbionts (61). It can also be hypothesized that combined expression of antimicrobial secondary metabolites may provide synergistic effects for egg defense.

Contrary to our initial hypothesis, no significantly differentially expressed genes were found between bacteria in the ANG and freshly laid eggs. During egg laying, the jelly coat material is secreted by the non-symbiotic nidamental glands and combined with bacteria-laden secretions of the ANG tubules (41). The chemical makeup of the egg jelly is unknown, but the similarity in bacterial gene expression suggests that the jelly may mimic the contents of the ANG tubules. Alternatively, ANG symbionts may be deposited with tubule contents, and thus are protected from extreme environmental changes. Sequencing of additional time points along the month-long egg development period and increased biological replicates could reveal more subtle changes in community gene expression over time.

Overall, the metatranscriptomes revealed expression of several pathways necessary for host-microbe interactions. In the *E. scolopes* light organ symbiosis, motility, chemotaxis, exopolysaccharide production, and quorum sensing have all been previously reported to be important for *V. fischeri* colonization fitness (62). In this study, we have found expression of all these pathways to varying levels among the ANG symbionts of mature, fully colonized squid. Some of these pathways, however, are not necessary for maintenance in the light organ symbiosis. For example, *V. fischeri* lose their flagella temporarily after light organ colonization (66). However, in *Loligo pealei* ∼10% of ANG bacteria were observed to be motile in adults (65). Further live imaging will be needed to confirm motility within *E. scolopes* ANG and eggs. Mucus secretion is also critical in recruitment of environmental *V. fischeri* to the light organ and found within the crypts where *V. fischeri* ultimately reside (63, 64), but utilization as a nutrient source is unclear. Mucosal epithelia may be the most common animal tissue involved in symbiotic associations (67), and expression of host mucin genes was detected in the ANG. Studies of mucin degradation have focused primarily on gut bacteria (68). The ANG symbiosis thus represents an exciting model to further examine the role of mucins in host-microbe interactions of defensive symbioses.

## Data Availability

All data is available under NCBI Genbank BioProject accession number PRJNA940485. Raw metatranscriptome reads are deposited in the NCBI Sequence Read Archive under accessions SRX19986921-SRX19986939 and raw metagenome reads under accession numbers SRX19972641-SRX19972644.

## Acknowledgements

We would like to thank Marcy Balunas for use of the freeze dryer, Bo Reese for sequencing technical assistance, and Artemis “Dyanna” Louyakis and the University of Connecticut Computational Biology Core for bioinformatic advice. We thank Allison Kerwin and Alecia Septer for helpful discussions during the preparation of this manuscript. Research was funded by National Science Foundation IOS 2247197 and 1557914 and the Gordon and Betty Moore Foundation 9349 and 12342 to SVN.

## Author Contributions

AMS and SVN conceptualized the study. AMS conducted experiments. AMS and SVN analyzed data, wrote, and edited the manuscript.

